# A Preclinical Alcohol BioBank: Samples from Behaviorally Characterized HS Rats for AUD Research

**DOI:** 10.1101/2025.04.30.651600

**Authors:** Michelle R. Doyle, Paola Campo, Selen Dirik, Maria G. Balaguer, Angelica R. Martinez, Marsida Kallupi, Giordano de Guglielmo

## Abstract

Alcohol use disorder (AUD) imposes a significant global health burden, yet effective treatments remain limited due to the scarcity of well-characterized biological sample repositories. To address this gap, we established the UCSD Alcohol BioBank, a comprehensive resource containing thousands of samples from over 700 genetically diverse heterogeneous stock (HS) rats. Modeled after successful cocaine and oxycodone biobanks, this repository utilizes the chronic intermittent ethanol vapor exposure (CIE) model, paired with oral self-administration, to characterize AUD-like behaviors, including ethanol consumption, preference, motivation, and withdrawal symptoms such as allodynia and anxiety-like behavior. Longitudinal samples (blood, urine, and feces) are collected before, during, and after ethanol exposure, while terminal samples (brain, heart, liver, kidneys, cecum, reproductive organs, adrenal glands, peripheral blood mononuclear cells) are obtained at intoxication, acute withdrawal, protracted abstinence, or from naive controls. Samples are preserved via snap-freezing or paraformaldehyde fixation to support diverse applications, including genomics, transcriptomics, proteomics, and neuroanatomy. The genetic diversity of HS rats enables genome-wide association studies (GWAS) to identify AUD-related genetic variants. Freely available to non-profit organizations at www.alcoholbiobank.org, with genetic and behavioral data deposited in public repositories, the Alcohol BioBank facilitates collaborative research to uncover biomarkers and develop novel therapies for AUD, addressing a critical need in addiction science.

## Introduction

Alcohol use disorders (AUDs) are pervasive in society, affecting an estimated 400 million people worldwide and contributing to approximately 2.6 million deaths annually (Organization, 2024). In the United States, approximately 29.5 million individuals aged 12 and older suffer from AUD, yet only about 7.6% receive any form of treatment (SAMHSA, 2023). Currently, there are three Food and Drug Administration (FDA)-approved pharmacological treatments for AUD— disulfiram, naltrexone, and acamprosate—but their efficacy is modest, and side effects often limit their use (Han et al., 2021). To improve current treatments and discover new ones, the individual differences in both the propensity to develop AUD-like behaviors and the response to treatment need to be better understood. A critical hurdle in identifying biomarkers and therapeutic targets is the limited number of repositories that include longitudinal samples with detailed characterization of AUD.

To address this gap, we have created the Alcohol BioBank (www.alcoholbiobank.org) which contains thousands of samples from over 700 individual heterogeneous stock (HS) rats (Hansen and Spuhler, 1984) and will continue to grow. The Alcohol BioBank, modeled after the successful cocaine and oxycodone biobanks (Carrette et al., 2021), aims to identify biomarkers for AUD and facilitate the development of novel therapies by leveraging the genetic diversity and detailed behavioral characterization of these rats. While preclinical models have identified potential biomarkers for AUD, translating these findings to humans has been challenging due to small sample sizes, lack of standardized protocols across laboratories, and the use of inbred strains that fail to capture human genetic diversity. Each of the over 700 rats in the biobank is fully characterized for AUD-like behaviors in a controlled environment using state-of-the-art oral self-administration models paired with chronic intermittent ethanol vapor exposure (CIE), a

well-established model of ethanol dependence (Roberts et al., 2000; Gilpin et al., 2008; Vendruscolo and Roberts, 2014; de Guglielmo et al., 2016; de Guglielmo et al., 2019; de Guglielmo et al., 2023; Doyle et al., 2024). Behavioral assessments include ethanol consumption, preference for ethanol over water, motivation under progressive ratio schedules, and withdrawal symptoms such as mechanical allodynia, anxiety-like behavior, and tolerance to the sedative effects of ethanol.

The HS rats, derived from crossbreeding eight genetically diverse founder strains (Hansen and Spuhler, 1984), allow for genome-wide association studies (GWAS) to identify genetic variants associated with AUD-related traits, supported by whole-genome sequencing of each individual (Chitre et al., 2020; Gunturkun et al., 2022; Lara et al., 2024; King et al., 2025; Kuhn et al., 2025). Longitudinal samples, including blood, urine, and feces, are collected throughout the behavioral experiments. Terminal samples, such as brain, heart, kidneys, liver, cecum, reproductive organs, and adrenal glands, are obtained at three key stages: during intoxication (13–15 hours of ethanol vapor exposure), acute withdrawal (7–10 hours post-vapor), and protracted abstinence (4 weeks post-vapor), as well as from naive rats. Samples are preserved using snap-freezing or paraformaldehyde fixation to ensure compatibility with various downstream applications.

The Alcohol BioBank is designed with an open-source mindset, making samples freely available to non-profit organizations at www.alcoholbiobank.org. Genetic and behavioral data will be deposited in public repositories such as the Rat Genome Database (Smith et al., 2020) or Gene Network (Sloan et al., 2016). This resource is particularly valuable for researchers outside the addiction field who lack the infrastructure for complex behavioral studies, enabling them to access well-characterized samples for multi-omics analyses and therapeutic testing.

Here, we introduce the Alcohol BioBank, providing methodological details and highlighting its potential applications to advance AUD research.

## Methods

### Subjects

Heterogeneous stock (HS) rats were provided by Leah Solberg Woods (Wake Forest University School of Medicine) and Abraham Palmer (University of California San Diego). The strain was originally created at NIH in the 1980s by crossbreeding 8 inbred strains (Hansen and Spuhler, 1984). In recent decades, the HS rat colonies have consisted of greater than 60 breeder pairs and used a breeding schematic for advanced intercrossed line that decreases inbreeding for future generations. In addition, only one male and one female offspring from each breeder pair were used in the behavioral characterization studies. At weaning, rats are implanted with a radio-frequency identification (RFID) chip to identify the individuals. The chip is scanned before every experiment to ensure the rat’s identity and make sure all data is properly linked to each rat, in order to ensure high fidelity data.

Rats were pair-housed in a 12h/12h reverse light-dark cycle (lights on at 20:00 during non-dependent phase or 21:00 during dependent phase). Rooms were temperature (20–22°C) and humidity (45–55%) controlled, and rats had with *ad libitum* access to tap water and chow (Envigo Teklad Rat Food Diet 8604) when not in an experimental session. With little exception (e.g., training sessions), experiments were generally conducted between 10:00 – 13:00. Subjects were handled on at least 3 occasions before beginning any experimental procedures.

### Drugs

A 10% v/v ethanol solution was made by mixing 95% ethanol with tap water and delivered at a volume of 0.1 ml/reinforcer. Quinine solutions were created by dissolving 0.1 g or 0.3 g quinine hydrochloride dihydrate (Sigma-Aldrich, Burlington, MA) in 1 L of 10% v/v ethanol solution. The water self-administer was tap water. For the loss of righting reflex task, a solution of 20% v/v ethanol was made by mixing 95% ethanol with sterile saline and injected intraperitoneally (i.p.) at a dose of 2.5 g of ethanol per kilogram of body weight.

### Mechanical nociception

Paw withdrawal thresholds were evaluated for each rat at baseline (before any ethanol exposure) and during acute withdrawal from ethanol dependence (**Fig 1**). Rats were allowed to habituate to the room for at least 30-min and to the small, clear chamber, which was placed on a metal grid for at least 10-min before measures were made. Paw withdrawal threshold was measured for each hind paw in triplicate using a dynamic plantar aesthesiometer for mechanical stimulation (electronic Von Frey, Ugo Basile). The filament was placed in the center of the hind paw and the force was automatically increased from 0 to 40g at a constant rate over 20-sec. At least 1 minute was allowed between measurements. The time and force at which the paw was retracted is recorded and averaged across the two paws and triplicate measures.

**Figure 1.**
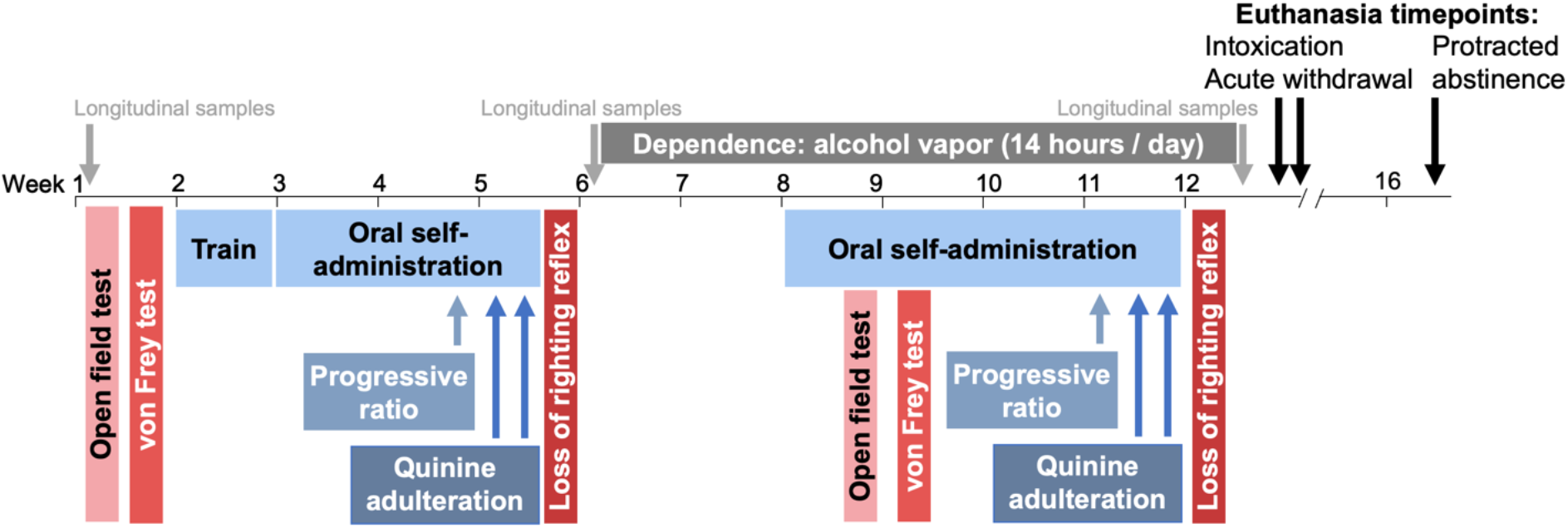
Experimental timeline where the time period where each behavioral assay is indicated. Alcohol consumption-related behaviors are in shades of blue and withdrawal-related behaviors are in shades of red. Longitudinal sample collection timepoints are indicated (before any alcohol exposure, after non-dependdent drinking, and during acute withdrawal in dependent rats). Rats were euthanized at the conclusion of the study, during one of three timepoints.

### Open field test

Entries made into the center of an open field were measured for each rat at baseline (before any ethanol exposure) and during acute withdrawal from ethanol dependence (**Fig 1**). Rats were allowed to habituate to the room for at least 30-minutes before testing began. Rats were placed into an open field (50 cm x 50 cm x 40 cm (height)) with black walls and gray floor and the center was defined as 16 cm x 16 cm area in the center of the open field. Behavior was tracked for 15-min using ANY-maze software (Wooddale, IL).

### Loss of righting reflex (LORR)

Duration of the loss of righting reflex after an injection of 2.5 g/kg ethanol (i.p.) was measured before induction of dependence (after pre-dependent self-administration) and after the conclusion of dependent self-administration (**Fig 1**). The times at which the rat was injected, lost its righting reflex (evaluated by placing rat on its back), and regained its righting reflex (evaluated by the rat righting itself three times within 15-sec) were recorded. The LORR duration (minutes that the rat was unable to right itself) at baseline was calculated as a measure of alcohol sensitivity. The change from the initial baseline measurement to that in acute withdrawal was considered a measure of ethanol tolerance.

### Self-administration procedure

#### Apparatus

Self-administration sessions were conducted in standard operant conditioning chambers (Med Associates, St. Albans, VT, USA). There were two retractable levers on the right wall, and a sipper cup containing two wells was located between the two levers. At the start of the session, the levers were inserted into the chamber and remained accessible for the entire duration of session. Two automated syringe drivers and 30-ml syringes were used to deliver the ethanol and water solutions into the right and left well, respectively, via Tygon tubing.

#### Training

To learn to self-administer oral solutions, rats were placed into operant chambers for a 16-hr self-administration session where tap water (0.1 ml/reinforcer) was delivered to a sipper cup upon completion of a fixed ratio (FR) 1 schedule of reinforcement (**Fig 1**). Two days later, rats were placed into operant chambers for a 16-hr self-administration session where ethanol (10% v/v in tap water; 0.1 ml/reinforcer) was delivered to a sipper cup the FR1. During both sessions, chow was available *ad libitum* but the only access to water or ethanol available was through responding on the active lever. Responses during the 0.6-sec timeout were recorded but had no programmed consequence.

#### Ethanol-water choice pre-dependence

After initial training, rats were allowed to self-administer ethanol (10% v/v) (right lever) or water (left lever) on FR1:FR1 schedule during daily 30-min sessions, where each response resulted in delivery of 0.1 ml of solution (**Fig 1**). After 9 daily (Monday-Friday) sessions, rats underwent a progressive ratio schedule test. During this, the session duration was increased to 45-min with a 15-min limited hold. Response requirement increased by 1 every other infusion for the first 20 infusions and then by 1 for every infusion after (1, 1, 2, 2, 3, 3, 4, 4, 5, 5, 6, 6, 7, 7, 8, 8, 9, 9, 10, 11, 12…), as previously described (Doyle et al., 2024). Response requirement increased independently on each lever. After the progressive ratio test, the next session was an ethanol-water choice session, followed by a “low quinine” session where rats could self-administer either 10% ethanol adulterated with 0.1 g/L quinine or water. Again, the next session was a baseline ethanol-water choice session, followed by a “high quinine” session where rats could self-administer either 10% ethanol adulterated with 0.3 g/L quinine or water. In total, the non-dependent self-administration phase lasted 14 sessions.

#### Self-administration during ethanol dependence

After rats were made physically dependent using the chronic, intermittent ethanol vapor exposure (CIE) model (see below for more details), self-administration sessions were resumed on Mondays, Wednesdays, and Fridays during acute withdrawal (7-10 hours after the vapor was turned off) (**Fig 1**). Rats self-administered ethanol and water under a similar procedure as before dependence, except, to allow time for escalation, 12 sessions were conducted before the PR and two quinine sessions were conducted. Again, there were intermediate ethanol-water choice sessions between the PR, low quinine, and high quinine sessions. In total, the dependent self-administration phase lasted 17 sessions.

#### Chronic intermittent ethanol vapor exposure to induce dependence

Rats were made physically dependent using the chronic, intermittent ethanol vapor exposure (CIE) model (Vendruscolo and Roberts, 2014; Kononoff et al., 2018; de Guglielmo et al., 2023; Doyle et al., 2024) (**Fig 1**). Rats were housed in ethanol vapor chambers (La Jolla Alcohol Research Inc, La Jolla, CA) and ethanol vapor was circulated for 14 hours/day. Ethanol vapor was slowly increased over the course of two weeks and BECs were measured twice per week until they reached an average of around 180 mg/dl, where most individual subjects were between 150-225 mg/dl.

#### Blood ethanol concentrations (BECs)

BECs were measured by collecting 0.1 ml of tail blood into a heparinized tube after pricking the tail vein with a 18G needle. Blood was spun in a centrifuge at 13000 RPM for 13 minutes to separate plasma. Plasma was then analyzed using gas chromatography, as previously described (de Guglielmo et al., 2023) or using an Analox AM1 Alcohol Analyser (Stourbridge, UK).

#### Euthanasia during intoxication, acute withdrawal, or protracted abstinence

At the end of the behavioral testing experiment, rats were euthanized during one of three timepoints: intoxication (after 13-15 hours of ethanol vapor), acute withdrawal (7-10 hours after the end of ethanol vapor exposure), or protracted abstinence (4 weeks after last ethanol vapor exposure)(**Fig 1**). For snap frozen samples, rats were euthanized using CO_2_ exposure and then decapitated. The brain was rapidly extracted and flash frozen using a 2-methylbutane and dry ice slurry. The other tissue was collected and frozen on dry ice. For fixed samples, rats were euthanized using CO_2_ exposure and then intracardially perfused with 150 ml of ice cold saline followed by 350 ml of 4% paraformaldehyde before tissue was collected. Fixed brains remained in paraformaldehyde at 4°C for 24-72 hours before cryoprotection in 30% sucrose with 0.1% sodium azide in phosphate buffered saline.

#### Tissue collected

For every experimental and naïve rat in the study, a variety of samples are collected longitudinally (throughout the experiment) or only at the terminal timepoint (**Table 1; Fig 1**). At the terminal timepoint, samples are collected when rats are intoxicated (after 13-15 hours of ethanol vapor), in acute withdrawal from alcohol (7-10 hours after the end of ethanol vapor exposure), or after a protracted abstinence period (4 weeks after last ethanol vapor exposure).

**Table 1.** List of samples that are collected longitudinally (left column) or only at the terminal timepoint (right column).

| **Longitudinal samples** | **Terminal samples** |
| --- | --- |
| Whole blood | Brain |
| Plasma | Heart |
| Serum | Liver |
| Urine | Kidney |
| Feces | Adrenal glands |
|  | Ovary or testes |
|  | Cecum |
|  | PMBCs |

#### Statistical analysis

For data shown in Fig 2 pre- and post-dependence, the days included were sessions 7-9 during the pre-dependence period and sessions 10-12 in the post-dependence period. These were the sessions immediately preceding the progressive ratio test. If rats received 3 or fewer rewards for all three sessions in the average, then preference was not calculated (n=4). If the injection for LORR was indicated as not accurate, data were excluded (n=48 of 380). If the ANY-maze software was unable to properly track the rat in the open field test, the data were excluded (n=8). Two samples were unable to be collected during the BAL time course (n=1 at timepoint 0 hours and n=1 at timepoint 8 hours).

**Figure 2.**
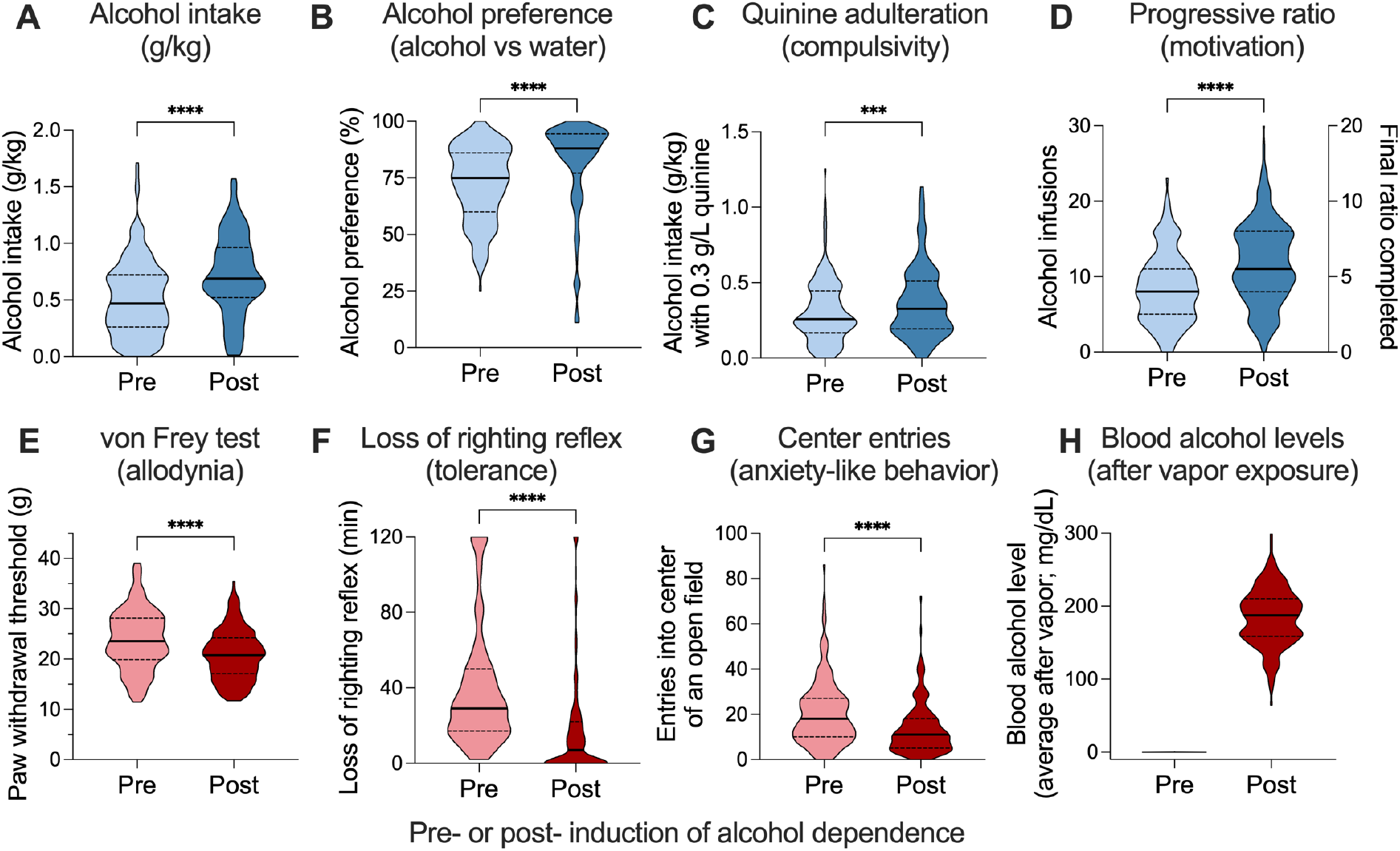
Comparisons between **A**. alcohol intake, **B**. alcohol preference, **C**. alcohol intake when 0.3 g/L quinine was added to the alcohol, and **D**. motivation to obtain alcohol using the progressive ratio schedule of reinforcement in rats pre- (light blue) or post- (dark blue) induction of alcohol dependence. Comparisons between **E**. paw withdrawal threshold as a measure of withdrawal-induced allodynia, **F**. loss of righting reflex as a measure of tolerance, and **G**. center entries in an open field as a measure of withdrawal-induced anxiety-like behavior pre- (light red) and post- (dark red) induction of alcohol dependence. **H**. Blood alcohol levels at the end of the 14 hours alcohol vapor exposure. In violin plots, thick lines indicate median and thin lines indicate quartiles. *** p < 0.001; **** p < 0.0001 for paired t-tests.

A paired t-test was used to analyze the pre- and post-dependence data in Fig 2. A one-way ANOVA was used to analyze the AUD-like phenotype data in Fig 3, with Sidak’s multiple comparison tests. When comparing change from baseline, a one sample t-test was also used to compare the values to 0 (Fig 3E-G). A mixed-effect analysis was used to analyze the BAL time course data in Fig 4. Area under the curve was calculated for each rat and plotted against the three-day average post-dependence intake and a linear regression was performed to determine the relationship.

**Figure 3.**
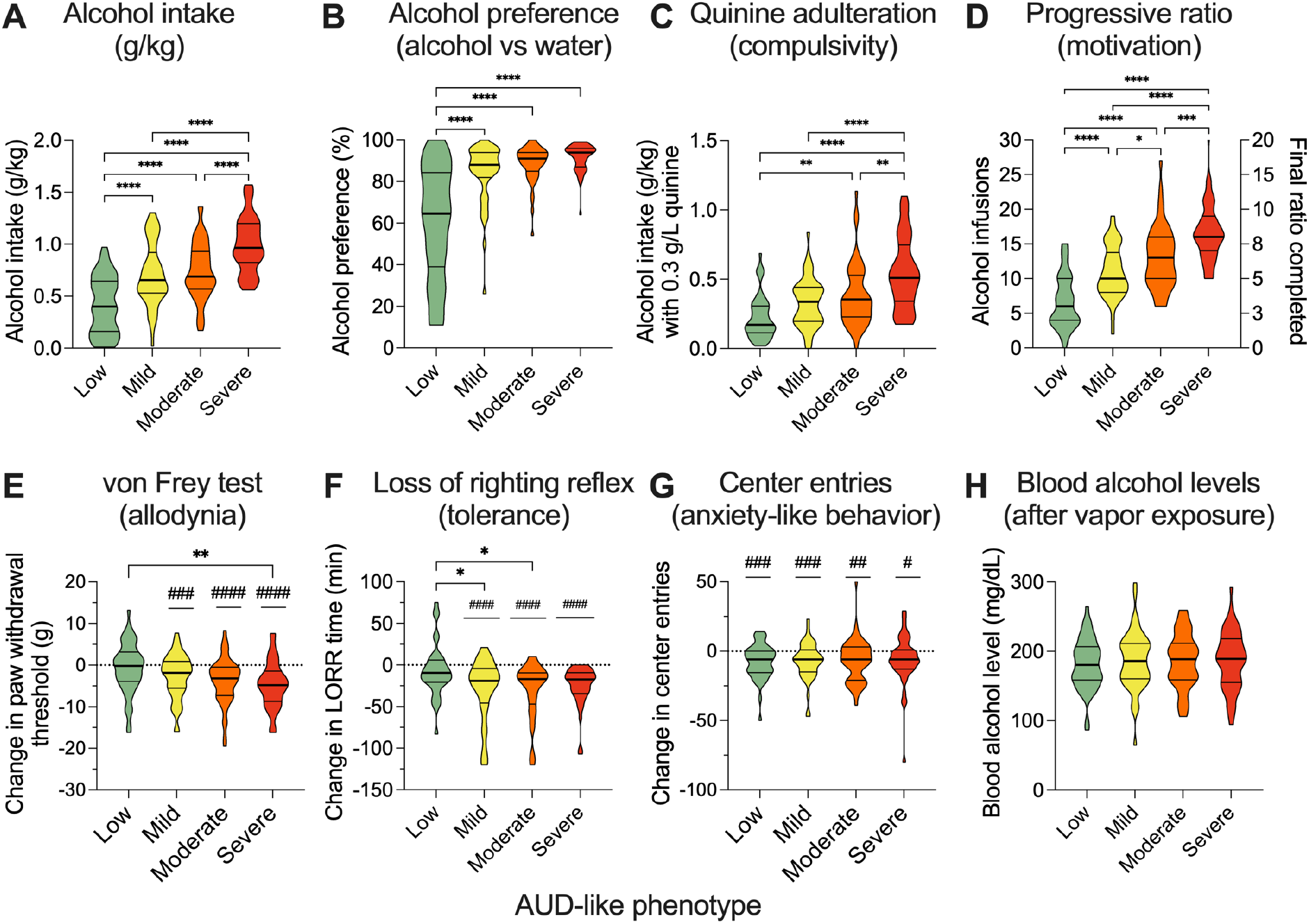
Comparisons between **A**. alcohol intake, **B**. alcohol preference, **C**. alcohol intake when 0.3 g/L quinine was added to the alcohol, and **D**. motivation to obtain alcohol using the progressive ratio schedule of reinforcement during dependence period in rats characterized as having a low (green), mild (yellow), moderate (orange) or severe (red) AUD-like phenotype. Comparisons in **E**. change in paw withdrawal threshold after dependence as a measure of withdrawal-induced allodynia, **F**. change in loss of righting reflex duration as a measure of tolerance, and **G**. change in center entries in an open field as a measure of anxiety-like behavior in rats with a low (green), mild (yellow), moderate (orange) or severe (red) AUD-like phenotype. **H**. Blood alcohol levels at the end of the 14 hours alcohol vapor exposure in rats within rats with a low (green), mild (yellow), moderate (orange) or severe (red) AUD-like phenotype. In violin plots, thick lines indicate median and thin lines indicate quartiles. * p < 0.05; ** p < 0.01; *** p < 0.001; **** p < 0.0001 for Sidak’s multiple comparison tests. # p < 0.05; ## p < 0.01; ### p < 0.001; #### p < 0.0001 versus 0 for one sample t-test.

**Figure 4.**
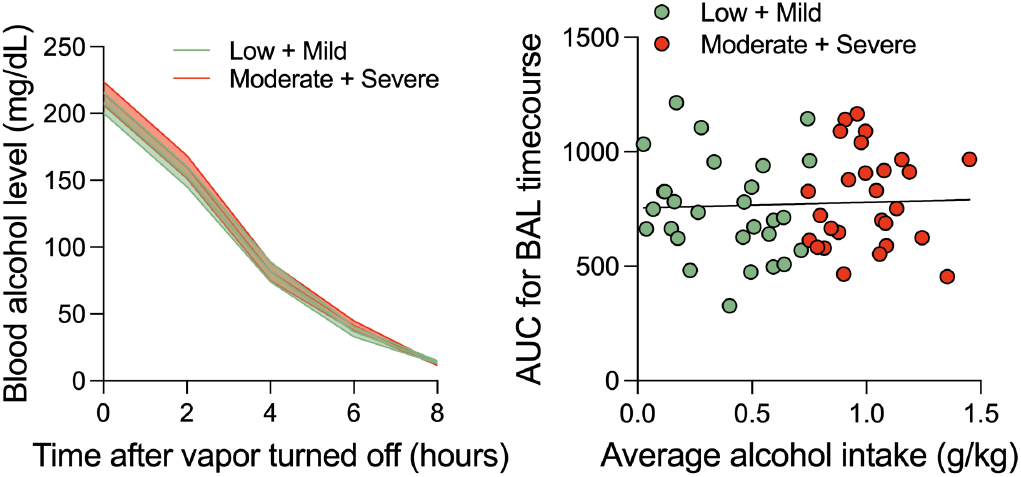
A. Time course of blood alcohol level (BAL) decay after the end of the alcohol vapor in rats that had a low or mild phenotype (green) versus moderate or severe phenotype (red). **B**. Lack of relationship between average alcohol intake during dependent self-administration and the AUC of the area under the curve (AUC) for the BAL time course.

To calculate AUD-like phenotype severity, we z-scored the measures of average alcohol intake during sessions 10-12 of dependence, reinforcers earned under progressive ratio, consumption of alcohol when 0.3 g/L of quinine was added, escalation of intake from pre- to post-dependence, tolerance in the LORR task, and allodynia in the von Frey test. Z-scores were calculated within sex and cohort to account for sex differences and potential cohort effects. The average of these 6 measures were used as an overall addiction index (Kallupi et al., 2020; de Guglielmo et al., 2024). The assignment to low, mild, moderate, or severe was based on dividing each cohort and sex into quartiles where the quartile with the lowest value was named low and the highest was named severe.

## Results

### Characterization of alcohol addiction-like behaviors

The timeline for the characterization of AUD-related behaviors is shown in Figure 1. We phenotype three cohorts of HS rats per year. Each cohort is composed by 96 HS rats (48 males and 48 females). We conduct von Frey (mechanical sensitivity) and open field tests (anxiety-like behavior) at baseline, before any alcohol self-administration and during acute withdrawal from alcohol dependence. We spend one week on training the rats to self-administer oral solutions (0.1 ml of 10% v/v ethanol or 0.1 ml of tap water), before undergoing the three-week self-administration protocol. Loss of righting reflex (LORR) is conducted at the end of the pre-dependence drinking phase to avoid exposing rats to a large, potentially aversive dose of ethanol before any drinking behavior. Over the course of two weeks, ethanol concentrations are increased in the ethanol vapor chambers. The rats remained receiving ethanol vapor for 14 hours/day throughout the rest of the experiment and conducted behavioral experiments (self-administration, open field, von Frey) during acute withdrawal, which was defined as 7-10 hours after ethanol vapor turned off. At the end of the self-administration studies, rats underwent a second LORR test to measure tolerance. Rats were then assigned to be euthanized during intoxication (euthanasia occurred while rats were in hour 13-15 of ethanol vapor exposure), acute withdrawal (7-10 hours after ethanol vapor turned off), or protracted abstinence (4 weeks after last ethanol vapor exposure).

Data presented in Figures 2–4 are derived from two cohorts of heterogeneous stock (HS) rats (n=190), providing a robust initial dataset from the UCSD Alcohol BioBank, which continues to expand with additional cohorts.

### Effects pre-and post-dependence using CIE

HS rats (n=190) went through the procedure as described and measured obtained during the pre-and post-dependence phases were evaluated (**Fig 2**). Rats self-administered more ethanol (t (189) = 8.280; p<0.0001; **Fig 2A**), had greater preference for ethanol (t (185) = 6.881; p<0.0001; **Fig 2B**), consume more ethanol when quinine is added (t (189) = 3.530; p=0.0005; **Fig 2C**), and showed more motivation to obtain ethanol under the progressive ratio schedule of reinforcement (t (189) = 8.013; p<0.0001; **Fig 2D**) after induction of dependence compared to their own pre-dependence baseline.

Paw withdrawal threshold in the von Frey test (t (189) = 7.161; p<0.0001; **Fig 2E**), time of loss of righting reflex as a measure of alcohol sensitivity (t (148) = 8.468; p<0.0001; **Fig 2F**), entries into the center of an open field as a measure of anxiety-like behavior (t (182) = 6.218; p<0.0001; **Fig 2G**) all decreased after induction of dependence compared to the pre-dependence baseline. Blood alcohol levels were measured after alcohol dependence and were, on average, 184.7 ± 2.83 (**Fig 2H**).

### Individual differences in phenotype severity

We calculate an addiction index for each of the rats and separated them into quartiles classified as low, mild, moderate, or severe, where rats with the lowest addiction index were considered low and those with the highest were severe. As expected, there were substantial individual differences and the groups differed significantly across all self-administration procedures, including alcohol intake (F (3, 187) = 35.56; p<0.0001; **Fig 3A**), alcohol preference (F (3, 186) = 37.14; p<0.0001; **Fig 3B**), alcohol consumption when adulterated with quinine (F (3, 186) = 19.14; p<0.0001; **Fig 3C**), and responding under a progressive ratio schedule (F (3, 186) = 50.71; p<0.0001; **Fig 3D**).

There was also a significant effect of group when evaluating withdrawal-induced allodynia (F (3, 185) = 4.208, p=0.0066; **Fig 3E**). All of the groups also showed significant allodynia (mild: t (47) = 3.511, p=0.001; moderate: t (46) = 4.677, p<0.0001; severe: t (45) = 5.681; p<0.0001)), except the low group (t (47) = 0.8823, p = 0.3821; **Fig 3E**). For the tolerance measure, there was a significant effect of group (F (3, 143) = 4.404, p=0.0054; **Fig 3F**). All of the groups also showed significant tolerance (mild: t (37) = 4.901, p<0.0001; moderate: t (32) = 5.141, p<0.0001; severe: t (439) = 6.745; p<0.0001)), except the low group (t (35) = 1.258, p = 0.2169; **Fig 3F**). There was no effect of group when evaluating center entries in an open field test as a measure of anxiety-like behavior (F (3, 178) = 0.08682; p=0.9672; **Fig 3G**); however, all groups showed significant decreases in center entries (low: t(44) = 3.983, p=0.0003; mild: t(46) = 3.688, p=0.0006; moderate: t(44) = 2.890, p=0.0060; severe: t(44) = 2.251, p=0.0295).

The behavioral differences between the groups were not mediated by group differences in blood alcohol levels generated by the CIE model (F (3, 187) = 0.2109; p=0.8887; **Fig 3H**).

### Blood alcohol level time course and relationship to self-administration

To confirm rats were in withdrawal during the behavioral experiments (conducted 7-10 hours after vapor turned off) and evaluate potential effects of ethanol metabolism on the behavioral assays, blood alcohol levels (BALs) were measured in a subset of rats (n=57) every two hours beginning from when the ethanol vapor ended until 8 hours later. There was a main effect of time (F (4, 210) = 585.0; p<0.0001), but no effect of phenotype of interaction (**Fig 4A**). There was no significant relationship between average ethanol intake in the dependence period and BAL time course AUC (R^2^ = 0.00019; p=0.752; **Fig 4B**).

## Discussion

The UCSD Alcohol BioBank constitutes a pivotal resource for advancing alcohol use disorder (AUD) research, complementing established cocaine and oxycodone biobanks (Carrette et al., 2021) at the University of California San Diego. Over the past five years, these repositories have distributed over 2,000 samples to 40 investigators across the United States and Europe, catalyzing significant advancements in addiction neuroscience (Carrette et al., 2022; Duttke et al., 2022; Kumaresan et al., 2023; Zhou et al., 2023; Okamoto et al., 2024; Vu et al., 2025). The Alcohol BioBank extends this paradigm, providing thousands of samples from over 700 genetically diverse heterogeneous stock (HS) rats, with data from two cohorts (n=190) reported herein. By facilitating comparative analyses across substances, the biobank enables elucidation of shared and distinct neurobiological mechanisms underlying alcohol, cocaine, and oxycodone addiction. Each sample is rigorously characterized with phenotypic and genotypic data, accompanied by comprehensive metadata, including animal identification, sex, age, body weight, sample weight, collection date, and preservation timestamps. Radio-frequency identification (RFID) chips and barcodes ensure precise traceability and transparency throughout the sample lifecycle.

Longitudinal samples (blood, urine, feces) and terminal samples (brain, heart, liver, kidneys, cecum, reproductive organs, adrenal glands) are collected at critical experimental timepoints: baseline, pre-dependence, dependence, intoxication, acute withdrawal, protracted abstinence, or from naive controls. Preservation via snap-freezing or paraformaldehyde fixation supports diverse applications, including genomics, transcriptomics, proteomics, metabolomics, and neuroanatomical studies. The chronic intermittent ethanol vapor exposure (CIE) model, a validated standard in AUD research (Roberts et al., 2000; Gilpin et al., 2008; Vendruscolo and Roberts, 2014; de Guglielmo et al., 2016; de Guglielmo et al., 2019; de Guglielmo et al., 2023; Doyle et al., 2024) underpins the biobank’s behavioral phenotyping, capturing escalation of ethanol consumption, compulsive drinking, and withdrawal symptoms such as allodynia, anxiety-like behavior, and tolerance. This multifaceted approach aligns with at least five DSM-5 criteria for AUD, enhancing translational relevance (Nieto et al., 2021).

Data from two cohorts (n=190) provide robust insights into AUD-like behaviors. Post-dependence, rats exhibited significant escalation in ethanol intake, heightened preference, sustained consumption despite quinine adulteration, and increased motivation under progressive ratio schedules (Fig. 2A–D), indicative of compulsive drinking and tolerance, hallmark features of AUD (Koob and Volkow, 2010). Withdrawal-induced manifestations, including mechanical allodynia, reduced loss of righting reflex duration, and diminished open-field center entries (Fig. 2E–G), underscore the negative affective and somatic states associated with dependence (Koob, 2022). These findings, enabled by the biobank’s standardized CIE protocol and comprehensive phenotyping, establish a robust platform for identifying biomarkers of compulsive drinking and withdrawal severity.

Individual differences in AUD-like phenotype severity (Fig. 3) further highlight the biobank’s utility. Rats classified as moderate or severe exhibited significantly elevated ethanol intake, preference, and compulsive drinking compared to low or mild groups (Fig. 3A–D), with corresponding increases in allodynia and tolerance (Fig. 3E–F). Notably, anxiety-like behavior was consistent across groups (Fig. 3G), suggesting a pervasive withdrawal effect amenable to molecular and neural dissection using biobank samples. The absence of a correlation between blood ethanol concentrations (BECs) and ethanol intake (Fig. 4B) indicates that behavioral variability is driven by neuroadaptive and genetic factors rather than metabolic differences, facilitating targeted investigations using biobank resources. These results support genome-wide association studies (GWAS) to identify genetic variants associated with AUD severity, leveraging the genetic diversity of HS rats and whole-genome sequencing (Hansen, 1984; Solberg Woods and Palmer, 2019).

The biobank’s samples are currently supporting cutting-edge investigations, as summarized in Table 2, including microbiome profiling to elucidate gut-brain interactions, metabolomics to identify AUD-related metabolic signatures, proteomics to uncover protein dysregulation, single-cell RNA sequencing to map cellular heterogeneity in brain Brain (Whole) regions such as the amygdala and nucleus accumbens, neuroanatomical studies to delineate circuit alterations, *in vivo* MRI to assess structural and functional brain changes, whole-brain imaging to visualize neural adaptations, and RNA methylation to investigate epigenetic modifications. These applications address critical questions regarding AUD susceptibility, progression, and therapeutic response, with potential to identify novel targets for mitigating compulsive drinking or withdrawal-induced allodynia.

**Table 2.** Ongoing collaborations utilizing of Alcohol BioBank samples in AUD research.

| Sample Type | Preservation Technique | Research Application |
| --- | --- | --- |
| Feces and Caecum | Snap-frozen | **Microbiome Profiling:** Studying gut-brain interactions in AUD-related behaviors. |
| Blood (Plasma, Serum) | Snap-frozen | **Metabolomics:** Identifying AUD-related metabolic signatures and biomarkers. |
| Brain (Various Regions) | Snap-frozen | **Proteomics:** Uncovering protein dysregulation in AUD pathways. |
| Brain (Amygdala, Nucleus Accumbens) | Snap-frozen | **Single-Cell RNA Sequencing:** Mapping cellular heterogeneity in AUD brain regions. |
| Brain (Various Regions) | Paraformaldehyde Fixation | **Neuroanatomical Studies:** Delineating circuit alterations in AUD. |
| Live Animals | None | ***in vivo* MRI:** Assessing structural and functional brain changes in AUD. |
| Brain (Whole) | Paraformaldehyde Fixation | **Whole-Brain Imaging:** Visualizing neural adaptations in AUD. |
| Brain (Amygdala) | Snap-frozen | **RNA Methylation:** Investigating epigenetic modifications in AUD-related brain regions. |

Human biobanks, such as the NIAAA’s COGA (1995) and UK Biobank (Sudlow et al., 2015), provide genotypic and phenotypic data for AUD research, with COGA including interviews and electrophysiological measures from 17,000 individuals (Dick et al., 2023). However, human biobank sample acquisition faces significant barriers (Coppola et al., 2019; Annaratone et al., 2021; Nieto et al., 2021). Ethical and legal requirements, including informed consent, IRB approvals, GDPR compliance, and pseudonymization, create complex governance and access delays (Harris et al., 2012; Coppola et al., 2019; Chandrashekar et al., 2022). Heterogeneous protocols hinder standardization, limiting data comparability (Coppola et al., 2019). Pre-analytical variability, such as post-mortem delays, degrades sample quality, impeding molecular analyses (Shabihkhani et al., 2014; Coppola et al., 2019; Annaratone et al., 2021). High costs and underuse, as seen in the EFS Centre-Atlantique biobank, challenge sustainability (Sapey et al., 2016; Annaratone et al., 2021). Self-reported data, prone to bias, reduce AUD phenotyping precision (Greenfield and Kerr, 2008; Nieto et al., 2021). Conversely, the preclinical Alcohol BioBank’s standardized CIE model, immediate sample collection, cost-effective open-access framework, and detailed phenotyping, including brain tissue access, provide high-fidelity data for AUD research. By integrating findings from preclinical and human biobanks, researchers can bridge translational gaps, leveraging the controlled, high-fidelity data from the Alcohol BioBank to complement the broader but less precise datasets from human repositories, thereby advancing the understanding and treatment of AUD.

Our behavioral protocol while robust, has its own limitations. The CIE model paired with measures of addiction like behaviors, do not fully recapitulate the human condition (Nieto et al., 2021). The HS rat model, despite genetic diversity, may not replicate human AUD’s sociocultural dimensions, requiring integration with human studies for translational relevance. Controlled experimental conditions, while minimizing variability, cannot fully account for complex environmental factors influencing human AUD (Koob and Volkow, 2010). These constraints highlight the need for complementary human research to validate preclinical findings.

The biobank is poised to advance AUD research through multi-omics integration, combining genomics, transcriptomics, proteomics, and metabolomics to elucidate molecular mechanisms. It supports preclinical testing of novel therapeutics, accelerating treatment development. Longitudinal studies can explore AUD progression and recovery, leveraging serial sample collections. The open-access model promotes global collaborations, enabling diverse investigations into addiction mechanisms (Carrette et al., 2021). Additionally, it serves as an educational resource, training emerging addiction researchers. These efforts will maximize the biobank’s impact on biomarker discovery and therapeutic innovation.

In summary, the Alcohol BioBank represents a pivotal resource for the AUD research community, offering unprecedented access to well-characterized biological samples from a genetically diverse rat population. Through its comprehensive behavioral phenotyping, rigorous sample preservation, and commitment to open science, the biobank is poised to drive transformative advancements in our understanding and management of AUD, addressing a pressing global health challenge.

## Author Contributions

GdG designed research; MRD, SD, AM, MB, MK performed research; GdG and MRD analyzed the data; MRD and GdG wrote the manuscript.

## References

(1995) The Collaborative Study on the Genetics of Alcoholism. Alcohol Health Res World 19:228–236.

Annaratone L, De Palma G, Bonizzi G, Sapino A, Botti G, Berrino E, Mannelli C, Arcella P, Di Martino S, Steffan A, Daidone MG, Canzonieri V, Parodi B, Paradiso AV, Barberis M, Marchio C, Alleanza Contro il Cancro P, Biobanking Working G (2021) Basic principles of biobanking: from biological samples to precision medicine for patients. Virchows Arch 479:233–246.

Carrette LLG, Corral C, Boomhower B, Brennan M, Crook C, Ortez C, Shankar K, Simpson S, Maturin L, Solberg Woods LC, Palmer AA, de Guglielmo G, George O (2022) Leptin Protects Against the Development and Expression of Cocaine Addiction-Like Behavior in Heterogeneous Stock Rats. Front Behav Neurosci 16:832899.

Carrette LLG et al. (2021) The Cocaine and Oxycodone Biobanks, Two Repositories from Genetically Diverse and Behaviorally Characterized Rats for the Study of Addiction. eNeuro 8.

Chandrashekar C, Shetty SS, Radhakrishnan R (2022) Evolution of biobanks and ethical governance for the emerging applications in biomedical research. J Oral Maxillofac Pathol 26:433–439.

Chitre AS et al. (2020) Genome-Wide Association Study in 3,173 Outbred Rats Identifies Multiple Loci for Body Weight, Adiposity, and Fasting Glucose. Obesity (Silver Spring) 28:1964–1973.

Coppola L, Cianflone A, Grimaldi AM, Incoronato M, Bevilacqua P, Messina F, Baselice S, Soricelli A, Mirabelli P, Salvatore M (2019) Biobanking in health care: evolution and future directions. J Transl Med 17:172.

de Guglielmo G, Simpson S, Kimbrough A, Conlisk D, Baker R, Cantor M, Kallupi M, George O (2023) Voluntary and forced exposure to ethanol vapor produces similar escalation of alcohol drinking but differential recruitment of brain regions related to stress, habit, and reward in male rats. Neuropharmacology 222:109309.

de Guglielmo G, Crawford E, Kim S, Vendruscolo LF, Hope BT, Brennan M, Cole M, Koob GF, George O (2016) Recruitment of a Neuronal Ensemble in the Central Nucleus of the Amygdala Is Required for Alcohol Dependence. The Journal of neuroscience : the official journal of the Society for Neuroscience 36:9446–9453.

de Guglielmo G, Kallupi M, Pomrenze MB, Crawford E, Simpson S, Schweitzer P, Koob GF, Messing RO, George O (2019) Inactivation of a CRF-dependent amygdalofugal pathway reverses addiction-like behaviors in alcohol-dependent rats. Nat Commun 10:1238.

de Guglielmo G et al. (2024) Large-scale characterization of cocaine addiction-like behaviors reveals that escalation of intake, aversion-resistant responding, and breaking-points are highly correlated measures of the same construct. Elife 12.

Dick DM, Balcke E, McCutcheon V, Francis M, Kuo S, Salvatore J, Meyers J, Bierut LJ, Schuckit M, Hesselbrock V, Edenberg HJ, Porjesz B, Collaborators C, Kuperman S, Kramer J, Bucholz K (2023) The collaborative study on the genetics of alcoholism: Sample and clinical data. Genes Brain Behav 22:e12860.

Doyle MR, Dirik S, Martinez AR, Hughes TE, Iyer MR, Sneddon EA, Seo H, Cohen SM, de Guglielmo G (2024) Catechol-O-Methyltransferase inhibition and alcohol use disorder: Evaluating the efficacy of tolcapone in ethanol-dependent rats. Neuropharmacology 242:109770.

Duttke SH, Montilla-Perez P, Chang MW, Li H, Chen H, Carrette LLG, de Guglielmo G, George O, Palmer AA, Benner C, Telese F (2022) Glucocorticoid Receptor-Regulated Enhancers Play a Central Role in the Gene Regulatory Networks Underlying Drug Addiction. Front Neurosci 16:858427.

Gilpin NW, Richardson HN, Cole M, Koob GF (2008) Vapor inhalation of alcohol in rats. Curr Protoc Neurosci Chapter 9:Unit 9 29.

Greenfield TK, Kerr WC (2008) Alcohol measurement methodology in epidemiology: recent advances and opportunities. Addiction 103:1082–1099.

Gunturkun MH, Wang T, Chitre AS, Garcia Martinez A, Holl K, St Pierre C, Bimschleger H, Gao J, Cheng R, Polesskaya O, Solberg Woods LC, Palmer AA, Chen H (2022) GenomeWide Association Study on Three Behaviors Tested in an Open Field in Heterogeneous Stock Rats Identifies Multiple Loci Implicated in Psychiatric Disorders. Front Psychiatry 13:790566.

Han B, Jones CM, Einstein EB, Powell PA, Compton WM (2021) Use of Medications for Alcohol Use Disorder in the US: Results From the 2019 National Survey on Drug Use and Health. JAMA Psychiatry 78:922–924.

Hansen C, Spuhler K (1984) Development of the National Institutes of Health genetically heterogeneous rat stock. Alcohol Clin Exp Res 8:477–479.

Harris JR et al. (2012) Toward a roadmap in global biobanking for health. European Journal of Human Genetics 20:1105–1111.

Kallupi M, Carrette LLG, Kononoff J, Solberg Woods LC, Palmer AA, Schweitzer P, George O, de Guglielmo G (2020) Nociceptin attenuates the escalation of oxycodone self-administration by normalizing CeA-GABA transmission in highly addicted rats. Proc Natl Acad Sci U S A 117:2140–2148.

King CP et al. (2025) Genetic Loci Influencing Cue-Reactivity in Heterogeneous Stock Rats. Genes Brain Behav 24:e70018.

Kononoff J, Kallupi M, Kimbrough A, Conlisk D, de Guglielmo G, George O (2018) Systemic and Intra-Habenular Activation of the Orphan G Protein-Coupled Receptor GPR139 Decreases Compulsive-Like Alcohol Drinking and Hyperalgesia in Alcohol-Dependent Rats. eNeuro 5.

Koob GF (2022) Anhedonia, Hyperkatifeia, and Negative Reinforcement in Substance Use Disorders. Curr Top Behav Neurosci 58:147–165.

Koob GF, Volkow ND (2010) Neurocircuitry of addiction. Neuropsychopharmacology 35:217–238.

Kuhn BN et al. (2025) Genome-wide association study reveals multiple loci for nociception and opioid consumption behaviors associated with heroin vulnerability in outbred rats. Mol Psychiatry.

Kumaresan V, Lim Y, Juneja P, Tipton AE, de Guglielmo G, Carrette LLG, Kallupi M, Maturin L, Liu Y, George O, Zhang H (2023) Abstinence from Escalation of Cocaine Intake Changes the microRNA Landscape in the Cortico-Accumbal Pathway. Biomedicines 11.

Lara MK, Chitre AS, Chen D, Johnson BB, Nguyen KM, Cohen KA, Muckadam SA, Lin B, Ziegler S, Beeson A, Sanches TM, Solberg Woods LC, Polesskaya O, Palmer AA, Mitchell SH (2024) Genome-wide association study of delay discounting in Heterogeneous Stock rats. Genes Brain Behav 23:e12909.

Nieto SJ, Grodin EN, Aguirre CG, Izquierdo A, Ray LA (2021) Translational opportunities in animal and human models to study alcohol use disorder. Transl Psychiatry 11:496.

Okamoto F et al. (2024) Y and mitochondrial chromosomes in the heterogeneous stock rat population. G3 (Bethesda) 14.

Organization WH (2024) Global status report on alcohol and health and treatment of substance use disorders. In: Global Report.

Roberts AJ, Heyser CJ, Cole M, Griffin P, Koob GF (2000) Excessive ethanol drinking following a history of dependence: animal model of allostasis. Neuropsychopharmacology 22:581–594.

SAMHSA CfBHSaQC (2023) 2023 National Survey on Drug Use and Health. In.

Sapey T, Py JY, Barnoux M, Tessier M, Dehaut F (2016) EFS Centre-Atlantique donor’s biobank: Ten years of samples usage. Transfus Clin Biol 23:95–97.

Shabihkhani M, Lucey GM, Wei B, Mareninov S, Lou JJ, Vinters HV, Singer EJ, Cloughesy TF, Yong WH (2014) The procurement, storage, and quality assurance of frozen blood and tissue biospecimens in pathology, biorepository, and biobank settings. Clin Biochem 47:258–266.

Sloan Z, Arends D, W. Broman K, Centeno A, Furlotte N, Nijveen H, Yan L, Zhou X, Williams R, Prins P (2016) GeneNetwork: framework for web-based genetics. J Open Source Softw 1:25.

Smith JR, Hayman GT, Wang SJ, Laulederkind SJF, Hoffman MJ, Kaldunski ML, Tutaj M, Thota J, Nalabolu HS, Ellanki SLR, Tutaj MA, De Pons JL, Kwitek AE, Dwinell MR, Shimoyama ME (2020) The Year of the Rat: The Rat Genome Database at 20: a multi-species knowledgebase and analysis platform. Nucleic Acids Res 48:D731–D742.

Sudlow C, Gallacher J, Allen N, Beral V, Burton P, Danesh J, Downey P, Elliott P, Green J, Landray M, Liu B, Matthews P, Ong G, Pell J, Silman A, Young A, Sprosen T, Peakman T, Collins R (2015) UK biobank: an open access resource for identifying the causes of a wide range of complex diseases of middle and old age. PLoS Med 12:e1001779.

Vendruscolo LF, Roberts AJ (2014) Operant alcohol self-administration in dependent rats: focus on the vapor model. Alcohol 48:277–286.

Vu T, Godbole S, Carrette LLG, Maturin L, George O, Saba LM, Kechris K (2025) Identification of Plasma Metabolites Responding to Oxycodone Exposure in Rats. Metabolites 15.

Zhou JL, de Guglielmo G, Ho AJ, Kallupi M, Pokhrel N, Li HR, Chitre AS, Munro D, Mohammadi P, Carrette LLG, George O, Palmer AA, McVicker G, Telese F (2023) Single-nucleus genomics in outbred rats with divergent cocaine addiction-like behaviors reveals changes in amygdala GABAergic inhibition. Nat Neurosci 26:1868–1879.

